# Fetal inflammatory signals regulate maternal investment in a pregnant marsupial

**DOI:** 10.1101/2025.08.16.669780

**Authors:** Daniel J. Stadtmauer, Jamie D. Maziarz, Oliver W. Griffith, Günter P. Wagner

## Abstract

Marsupial pregnancy is strikingly short: placental attachment in the gray short-tailed opossum *Monodelphis domestica* lasts only two days. This brevity has been attributed to a maternal immune response to fetal contact which only eutherian mammals have evolved mechanisms to tolerate. The attachment period is characterized by a spike in inflammatory signaling, development of an expanded uterine capillary network, and exponential fetal growth. Several inflammatory cytokines, including interleukin-1A (IL-1A) and interleukin-6 (IL-6), are produced primarily by fetal cells. We hypothesized that placental cytokines function as solicitation signals that increase maternal investment. To test this, we treated pregnant opossums with inhibitors of IL-1 and IL-6 during the rapid growth phase. Inhibition of IL-1 and IL-6 signaling significantly increased average biomass per fetus (+12% and +10%), and as such these signals impose costs, rather than direct benefits, to intrauterine growth. However, uninhibited controls showed greater surviving litter sizes than IL-1-inhibited animals, suggesting that IL-1A promotes offspring survival. Single-cell transcriptomes reveal that maternal vascular endothelial cells, perivascular cells, and fibroblasts are the primary targets of fetal IL-1A, and maternal cells simultaneously upregulate antagonists IL1R2 and IL1RN, suggesting resistance to fetal signaling. Placental transcriptomics reveals that the cytokine surge is restricted to the final day of pregnancy when placental cells fuse to form syncytial knots, and that these cells produce additional vasomodulatory signals including a truncated isoform of VEGFA. Maternal cells, in contrast, increase production of functional antagonists *IL1R2* and *IL1RN*, suggesting resistance to fetal signals. We propose that marsupials co-opted inflammatory signals to perform a novel function promoting fetal survival through maternal vascular remodeling.

## Introduction

The life history dichotomy between eutherian (placental) mammals and marsupials has long puzzled biologists. Marsupials give birth to highly altricial neonates which must complete development attached to the nipples, whereas eutherian mammals have evolved extended gestation which can produce precocial offspring. Major macroevolutionary trends, such as the extinction of many South American marsupials in the late Cenozoic upon introduction of eutherians to the continent, have been attributed to these differences in reproductive biology (Cox, 1977; Lillegraven, 1975). Marsupial short gestation has been attributed to a maternal immune response to paternally-derived fetal antigens which only eutherian mammals have evolved mechanisms to tolerate (Lillegraven, 1975). Under this hypothesis, marsupials are developmentally constrained from evolving extended gestation. Alternatively, it has been proposed that marsupials have short gestation instead due to selection for lactation rather than lack of developmental opportunity to evolve extended gestation (Hayssen et al., 1985; Renfree, 1983, 1981), and thus the placentation stage is shortened in order to proceed quickly to lactation.

Transcriptomic studies have lent apparent support to the immunological constraint hypothesis by revealing a pronounced production of pro-inflammatory cytokines at the utero-placental interface following dissolution of the shell coat and uterine attachment in the gray short-tailed opossum (Griffith et al., 2017; Hansen et al., 2017) and the fat-tailed dunnart (Dudley et al., 2024). However, two elements not predicted in the original immunological constraint model have also been discovered. First, the attachment response is predominantly driven by innate immunity - specifically inflammation - rather than adaptive immunity (Chavan et al., 2017; Griffith et al., 2017). Second, many cytokines have been discovered to be produced by the fetus via placental trophoblast cells, contrary to an antigen- or damage-induced inflammatory response which would presumably be maternally driven (Chavan et al., 2021; Stadtmauer et al., 2025; summarized in Stadtmauer and Wagner, 2020).

Beyond defense against non-self invaders, inflammation induces local edema and angiogenesis (Ashley et al., 2012). These sequelae of inflammation hold a latent potential for evolutionary co-option to facilitate nutrient delivery. This potential has been realized in multiple contexts, including the origin of mammalian lactation, where NF-κB and antibacterial enzymes have acquired secondary roles roles regulating resource allocation (Vorbach et al., 2006) as well as an evolutionarily novel brooding tissue in ricefish whose development depends upon inflammatory processes (Hilgers et al., 2022). Previously, we speculated that marsupials may have leveraged these vasomodulatory effects to enhance nutrient provisioning during placentation (Stadtmauer and Wagner, 2020). The repeated co-option of inflammation in evolutionarily novel modes of offspring provisioning raises the question: *does inflammation in marsupial pregnancy benefit the fetus*?

In the gray short-tailed opossum *Monodelphis domestica*, trophoblast cells fuse during the last 2 days of its 14.5-day gestation to form multinuclear aggregates known as syncytial knots (Mate et al., 1994; Zeller and Freyer, 2001). These cells produce *IL1A*, *IL6*, and other cytokines, and their development coincides with rapid growth of the fetus and glandular opening and the development of a subepithelial capillary network in the endometrium (**Figure 1**) (Griffith et al., 2019; Harder et al., 1993; Rose, 1989; Stadtmauer et al., 2025). Here, by identifying the cellular targets of placental inflammatory cytokines and pharmacologically inhibiting them to identify their functional consequences, we test whether fetal-driven inflammation directly influences maternal investment.

**Figure 1.**
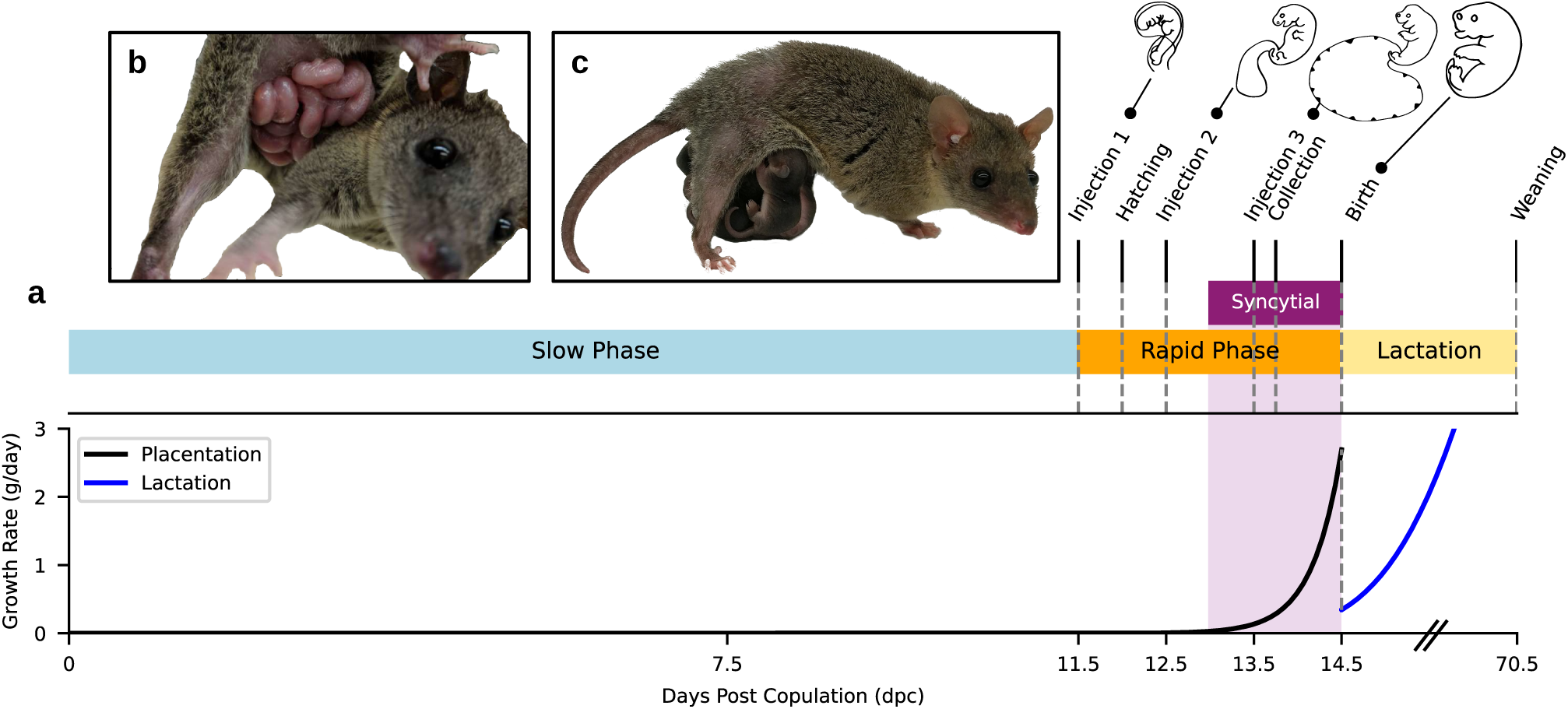
Pregnancy timeline in the opossum. **a**. Timeline of opossum development, showing the slow vs. rapid developmental phases and the onset of syncytial knot formation. Shell coat hatching occurs at 12 dpc and syncytialization on day 13. Lactational period from 14.5-70.5 dpc is linearly compressed for clarity. The approximate rate of biomass acquisition (g/day) is estimated below; placental growth is assumed to occur exponentially (Rose, 1989) and lactational growth via a Gompertz function (Dove and Cork, 1989) from measurements detailed in Table S1. Timing of anti-inflammatory injection experiments are marked. Illustrations are drawn after art by Lynne Selwood. **b.** *M. domestica* litter on the day of birth. ©Oliver Griffith. **c.** Litter ∼1 month into lactation period. ©Daniel Stadtmauer.

## Results

### Cellular sources and targets of placental cytokines

First, we identified fetal-maternal signals with the necessary temporal and cell type specificity required to be potential inflammatory solicitation signals. We compared bulk placental gene expression between day 13.5 and day 12.5 of gestation to isolate changes to the fetal ligand repertoire after syncytial knot formation on day 13, and re-analyzed single-cell transcriptomes from the day 13.5 dpc interface (Stadtmauer et al., 2025) and to identify endometrial target cells of fetal cytokines. Then, we tested the effects of inhibiting three cytokines – PGE_2_, IL-6, and IL-1 – on litter size and fetal biomass in order to gain insights into the functional significance of these signals.

#### Placental cytokine production coincides with syncytial knot development

The full set of genes with both placental specificity and temporal acuity to late pregnancy was identified by bulk transcriptomic comparison. 850 genes showed moderate or greater expression (>30 transcripts per million, TPM) in the 13.5 dpc placenta, but not in the 13.5 dpc-equivalent pseudopregnant endometrium or the 12.5 dpc placenta (**Figure 2a**). These included *IL1A, IL6, PTGES*, *VEGFA, IL10* and *IL17A* (**Figure 2a-b**). *HAND1*, a general trophoblast marker, was expressed in the placenta at both 12.5 and 13.5 dpc, while *GCM1*, a syncytial marker, appeared only at 13.5 dpc, consistent with syncytial knot formation after 13 dpc (**Figure 2a-b**). Together, this pattern suggests that inflammatory cytokine production begins after 12.5 dpc, coincidental with the shedding of egg coverings, and is located in the syncytial knots.

**Figure 2.**
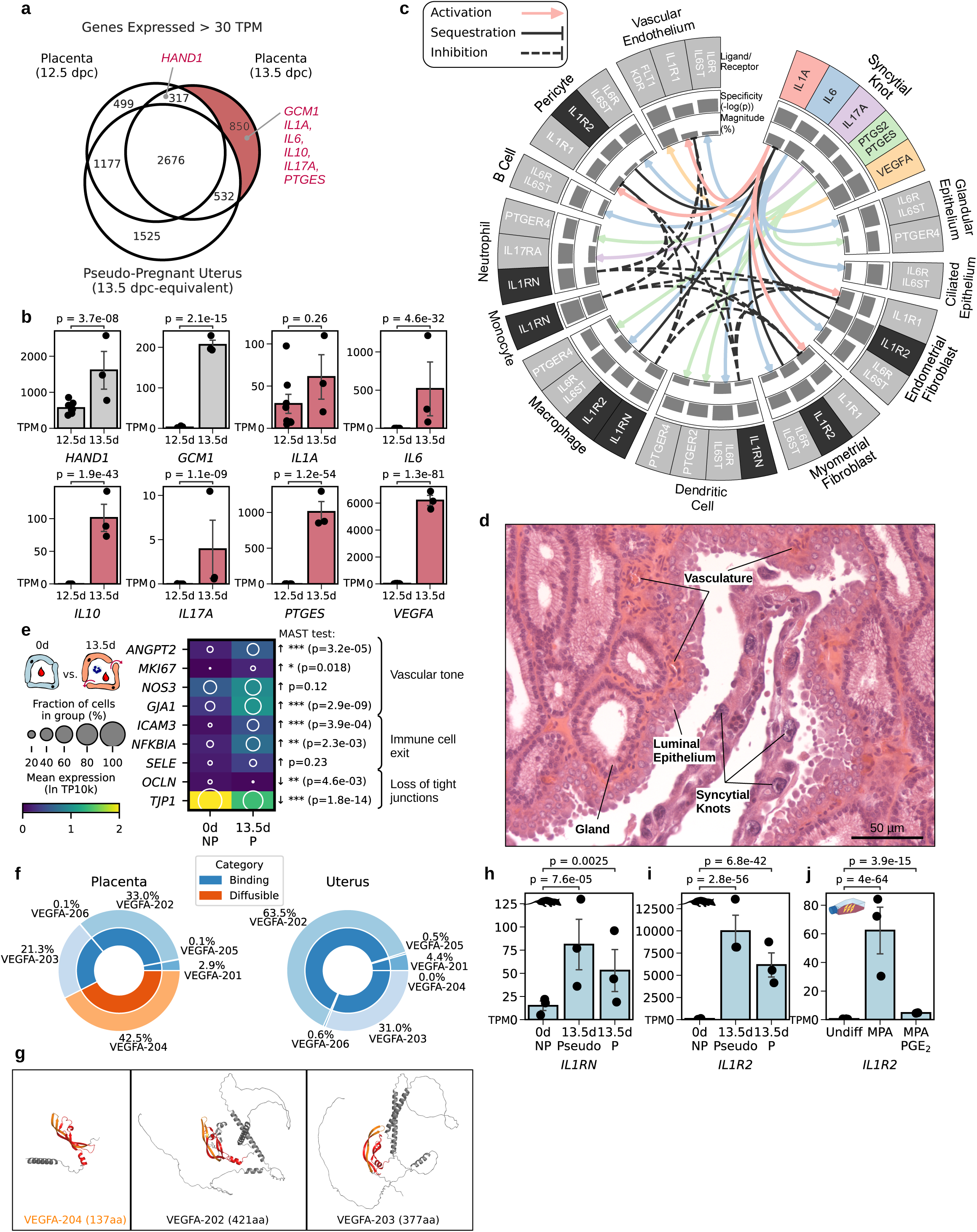
Temporal and cell type specificity of inflammatory ligands, receptors, and physiological inhibitors in the last day of opossum pregnancy. **a**. Venn diagram showing genes passing a moderate expression threshold (>30 TPM) in 13.5-day placental membranes (after syncytial knot formation), 12.5-day placental membranes (before syncytial knot formation), and 13.5-day-equivalent pseudo-pregnant uterus. **b.** Expression of select trophoblast cell type markers and inflammatory mediators before (12.5d) and after (13.5d) syncytialization from the same data sets as (**a**). **c.** Fetal-maternal signaling via inflammatory mediators in normal opossum pregnancy inferred from single-cell transcriptomes from the 13.5 dpc utero-placental interface (Stadtmauer et al., 2025). Magnitude barplots reflect the percent of cells in the cell type cluster with nonzero expression of the ligand or receptor gene (or lesser of the two for pairs like PTGS2+PTGES). Specificity barplots reflect LIANA+ robust rank aggregate p-value scores. Only interactions exceeding expression levels of 40 counts per million and 15% of cells in the cluster for ligand and receptor are shown. **d.** Hematoxlin and eosin-stained histological section showing apposition of syncytial knots to uterine lumen and underlying capillary networks at 13.5 dpc. Re-imaged in expanded view from supplementary figures of (Stadtmauer et al., 2025). **e.** Expression of markers of vascular permeability to nutrients *(ANGPT2, NOS3, GJA1),* vascular proliferation *(MKI67),* leukocyte migration *(ICAM3, SELE)*, cytokine response *(NFKBIA)*, and tight junction integrity *(OCLN, TJP1)* in vascular endothelial cell single-cell transcriptomes from non-pregnant and 13.5 dpc uteri. **i.** Relative *VEGFA* isoform abundances in 13.5 dpc placenta versus 13.5 dpc uterus show fetal-specific expression of the truncated splice variant *VEGFA-204*, equivalent to human VEGF_111_. **g.** Predicted protein stuctures (Jumper et al., 2021) of VEGFA-204 compared to co-expressed VEGFA-202 and VEGFA-203 show conservation the receptor-binding domain encoded by exons 2 and 3 (red and orange) but differ in the presence or absence of additional heparin and neuropilin-binding domains encoded by exons 5-7. **h-i.** Expression of *IL1RN* (**h**) and *IL1R2* (**i**) in bulk transcriptomes from non-pregnant (0d NP), 13.5 dpc-equivaelent pseudo-pregnant (13.5d Pseudo), and 13.5 dpc pregnant (13.5d P) individuals. **j.** Expression of *IL1R2* in *in vitro* cultured opossum endometrial stromal fibroblasts in response to 2 days treatment with either 1 µm of progesterone analog medroxyprogesterone 17-acetate (MPA), 1 µm MPA plus 10 µm PGE_2_ (MPA/PGE_2_) or growth medium-only control (Undiff). P-values in (**b**) and (**h-j**) are Benjamini-Hochberg-corrected Wald test results from pyDESeq2 on read counts, whereas p-values in (**e**) are likelihood ratio test p-values from MAST.

#### Maternal targets include glands and vasculature

We re-analyzed single-cell transcriptomes from the day 13.5 fetal-maternal interface (Stadtmauer et al., 2025) to identify maternal targets of placental cytokines. The IL-1A receptor *IL1R1* was largely absent from maternal immune cells, but expressed by vascular endothelial cells, pericytes, and endometrial stromal fibroblasts (**Figure 2c**). The IL-6 receptor *IL6R* and its co-receptor *IL6ST* were expressed in both immune (macrophages, dendritic cells, and B cells) and non-immune cells (glandular and ciliated epithelial cells, smooth muscle cells, and fibroblasts) (**Figure 2c**). The PGE_2_ receptor PTGER4 was also expressed in immune cells. Of these targets, vascular endothelial cells and glandular epithelial cells provide the most direct routes to affect nutrient allocation. As such, we compared endothelial cell type-specific transcriptomes in 13.5 dpc uterus to non-pregnant samples to investigate possible consequences of this signaling.

#### Fetal signaling alters vascular permeability and tone

The opossum placenta is separated from maternal vascular endothelial cells and their surrounding pericytes by a thin uterine epithelium, a short distance which leaves open the potential for paracrine signaling (**Figure 2d**). Maternal endothelial cells at 13.5 dpc expressed higher levels of vasodilation markers *ANGPT2* and *NOS3* (Balligand et al., 2009; Dai et al., 2021), proliferation marker *MKI67*, permeability marker *GJA1*, and leukocyte adhesion molecules (*ICAM3*, *SELE*), whereas tight junction markers *OCLN* and *TJP1* were downregulated (**Figure 2e**). These changes coincide with the histological development of a subepithelial capillary network and with the regression of sub-epithelial fibroblasts (Griffith et al., 2019). Intriguingly, *IL1R1* is expressed in both the cells of the vasculature (endothelial cells and pericytes) and fibroblasts, consistent with a role of IL-1 in both components of this late-gestation endometrial remodeling.

#### Syncytial knots produce a diffusible VEGFA isoform

*VEGFA* is abundantly expressed (>6000 transcripts per million; TPM) in the 13.5 dpc opossum placenta (**Figure 2b**), and ranked among the top 10 most abundantly transcribed genes by syncytial knot cells. Isoform-level quantification showed that 42.5% of placental *VEGFA* transcripts lacked exons 5-7 (**Figure 2f**), encoding a truncated peptide (VEGFA-204) analogous to human VEGF_111_, which is more distantly diffusible and potently angiogenic due to the loss of heparin and neuropilin binding sites and protease cleavage motifs (**Figure 2g**) (Mineur et al., 2007). This isoform was unexpressed in maternal tissues. Its rise to high levels coincides with rapid vascular expansion during late gestation, and is consistent with a primary action on maternal vascular development.

#### Maternal cells express IL-1 antagonists

Maternal immune cells expressed the competitive IL-1 receptor inhibitor *IL1RN*, but not the true receptor *IL1R1* (**Figure 2c,h**). *IL1R2*, a decoy receptor that sequesters IL-1A, was highly expressed by fibroblasts (>9000 TPM) (**Figure 2c**) and upregulated in abundance specifically at late gestation (**Figure 2i**), suggesting that it is not a constitutive product but instead an evolved regulatory response. *In vitro,* opossum endometrial stromal fibroblast expression of *IL1R2* is progesterone-responsive (**Figure 2j**), and pseudo-pregnant animals express high levels as well (**Figure 2i**). These results suggest that maternal cells actively buffer fetal IL-1A.

In sum, transcriptomic analyses demonstrate that receptors for placental cytokines, in particular IL-1A, are indeed expressed in maternal cell types which could plausibly have effects on nutrient allocation and resorption rate, and that maternal cells may exhibit a dynamic counter-regulation of fetal inflammatory signals.

### Functional inhibition of placental inflammation

Pregnant females received daily subcutaneous injections on 11.5, 12.5, and 13.5 dpc with either saline (n = 8), meloxicam (n = 6), anakinra (n = 7), or tocilizumab (n = 5), with tissue collected at 13.75 dpc. Litter size and total fetal biomass were recorded to estimate maternal nutrient allocation. Two experiments (1 saline, 1 tocilizumab) lacked biomass data due to damage during sample collection.

#### PGE_2_ inhibition does not affect fetal allocation

We previously measured uterine PGE_2_ levels in opossums and found a ∼20 fold increase during late gestation (Erkenbrack et al., 2018). We attempted to block this increase with 1 mg daily injections of meloxicam, an inhibitor of the prostaglandin H_2_ synthase PTGS2. However, 1 mg daily injections failed to reduce uterine PGE_2_ levels as measured 6 hours following the last dose (45 ng/mg protein vs. 55 ng/mg, *p* = 0.66) (**Figure S1a**) or to alter fetal biomass. An injection protocol delaying each injection by 5 hrs, with collection 1 hr following the last dose, reduced PGE_2_ to 9 ng/mg protein (*p* = 1.0 × 10^−6^) (**Figure S1a**), showing that meloxicam is effective in the opossum but its effects on PGE_2_ are short-lived, likely due to rapid drug metabolism. This short-term decrease in PGE_2_ levels was still without effect on biomass or litter size (**Figure 3b-c**). Continuous delivery via osmotic pump (from 12.75-13.75 dpc) also lowered PGE_2_ (11 ng/mg; **Figure S1a**), but had no measurable effect on biomass or litter size (**Figure S1b**), suggesting that transient depletion of prostaglandins is insufficient to alter nutrient allocation. We conclude that meloxicam is not ineffective in the opossum, but the lack of effect due to rapid restoration of PGE_2_ levels between 1 and 5 hours after injection prevented robust inhibition.

**Figure 3.**
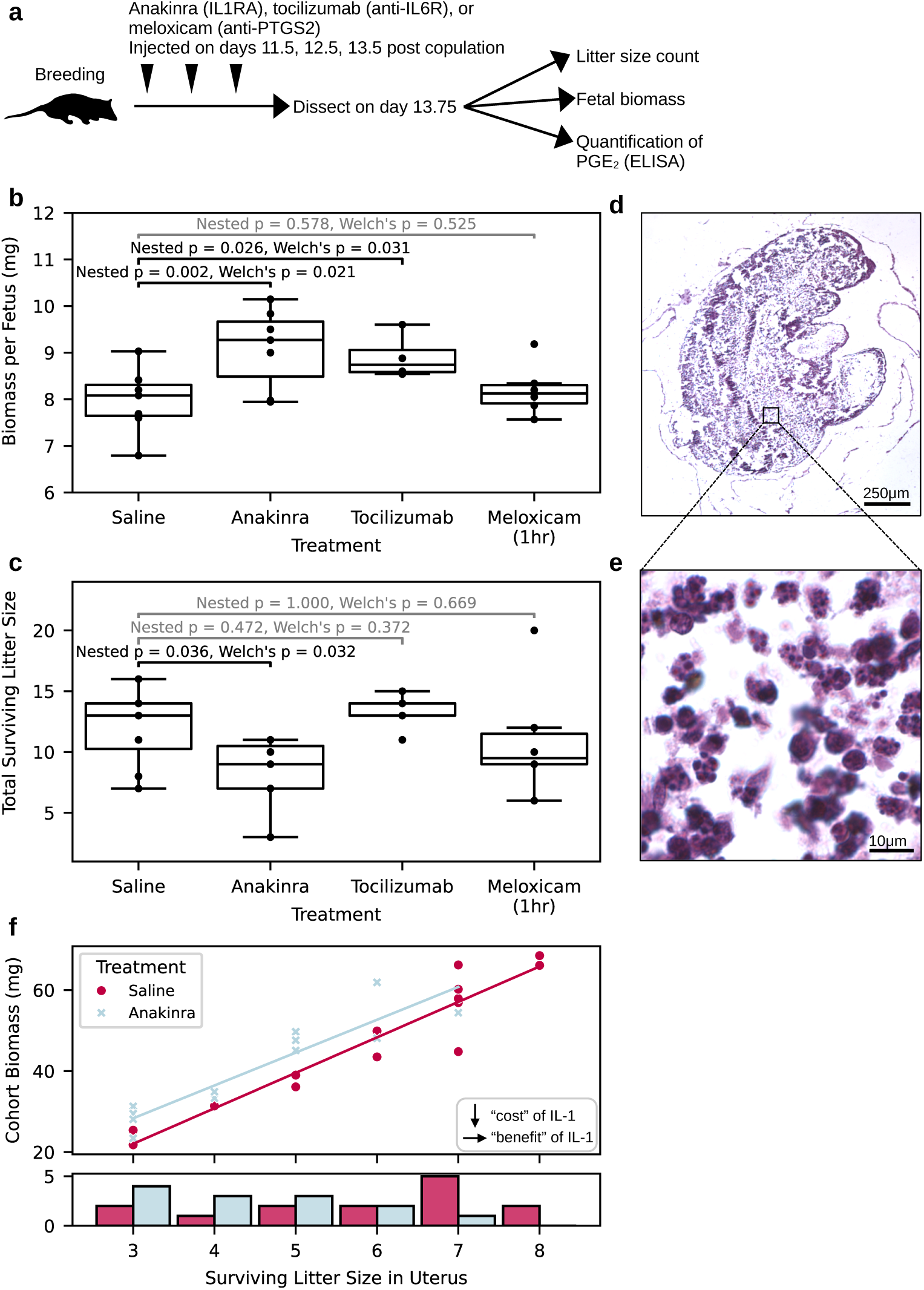
Anti-inflammatory intervention into opossum pregnancy. **a.** Experimental design. **b.** Effect of anakinra, tocilizumab, and meloxicam on fetal biomass. **c.** Effect of anakinra, tocilizumab, and meloxicam on litter size. **d.** Hematoxylin and eosin histology of resorbed embryo remnants. **e.** Zoomed panel from (**d**) showing resorbed matter cleanup by granulocytes and lymphocytes. **f.** Relationship of total fetal biomass within a uterus (cohort biomass) to number of fetuses inside (surviving litter size in uterus) in saline and anakinra treatment groups shows a downward shift in the uninhibited control group. Histogram below shows the number of observations within each treatment group, following a trend towards larger surviving litter size in the saline group.

#### IL-6 blockade increases fetal biomass acquisition

Functional effects of IL-6 were examined using tocilizumab, a monoclonal antibody which competitively binds to the IL-6 receptor. Tocilizumab-treated animals showed a significant (two-tailed Welch’s *p* = 3.1 × 10^−2^, mixed linear model *p* = 2.6 × 10^−2^) increase in fetal dry mass per fetus of 10% (from 7.97 to 8.90 mg, *d* = 1.54) (**Figure 3b**), but no significant change in litter size (12.0 vs. 13.2, *p* = 0.372) (**Figure 3c**). This caused a non-significant trend in the opposite direction as predicted, towards greater total maternal investment (+20 mg, *p* = 0.127) with IL-6 suppression.

#### IL-1 inhibition leads to fewer, larger surviving offspring

Effects of fetal IL-1A were investigated using anakinra, modified recombinant IL1RN which antagonizes the IL-1 receptor, effectively supplementing existing maternal IL1RN at extreme levels. Anakinra-treated animals showed a significant (two-tailed Welch’s *p* = 2.1 × 10^−2^) increase in fetal dry mass of 12% (from 7.97 to 9.10 mg, *d* = 1.43) (**Figure 3b**). This increase was coupled with a significant decrease in average litter size by 3.7 (from 12.0 to 8.3, *d* = −1.24, two-tailed Welch’s *p* = 3.2 × 10^−2^) (**Figure 3c**).

Traces of actively resorbed embryos were occasionally observed in both untreated and treated uteri. These took the form of 1-2 mm white masses in the shape of fetuses (**Figure 3d**) nearly entirely consisting of granulocytes (**Figure 3e**), consistent with descriptions of uterine resorption in rodents (Drews et al., 2020). The number of actively resorbing embryos observed (9 across all animals) was insufficient to to statistically determine frequency under different treatment conditions, but nevertheless suggest a plausible mechanism to affect surviving litter size.

One possible determinant of fetal biomass is a passive scaling effect: if space within the uterus is limited, the size and number of offspring could be nonlinear and governed by a trade-off, with smaller litters having larger fetuses and diminished returns on fetal size in larger litters due to crowding. If so, IL-1 could conceivably affect survival alone, with growth effects merely a byproduct of reduced crowding. However, cohort biomass showed a linear increase with litter size in each uterus (**Figure 3f**), suggesting that no such effect was at play. This rules out passive scaling as the sole driver.

## Discussion

Our findings suggest that inflammatory signals in *M. domestica* play complex roles in shaping maternal investment. Cytokines such as IL-1A, IL-6, IL-10, IL-17A and PGE_2_ are traditionally associated with immune activation, yet at the opossum fetal-maternal interface, their receptors are primarily expressed on non-immune maternal targets such as glandular epithelium, vascular endothelium, and stromal cells. Functional experiments on two of these pathways, IL-1 and IL-6, demonstrate that neither has the properties expected of simple solicitation signals positively associated with fetal growth; inhibition of both did not reduce fetal biomass, but rather increased it. These signals appear to act beyond direct nutrient and immune regulation to regulate the development of a capillary network which enables fetal survival.

Once the shell coat degrades at 12 dpc, fetuses secure placental territory and receive oxygen and nutrients from endometrial glands and vasculature. One day later, the final, syncytial stage of placentation begins with the differentiation of giant syncytial knot cells (**Figure 1a**). Fetal syncytial knot cells are highly angiogenic, producing the highly-potent truncated *VEGFA* isoform (VEGF_111_) and high levels of inflammatory cytokines. This appears to initiate the development of a subepithelial endometrial capillary network, whose development shows signs of endothelial cell permeability by loss of tight junctions and increased vascular tone (**Figure 2d-e**), enabling exponential fetal growth (**Figure 1a**). That capillary network development depends upon fetally-derived signals has been previously shown by observations that it is absent in pseudopregnant animals (Griffith et al., 2019). Here, our identification of *IL1R1* expression in endothelial and perivascular cells, and finding of a functional link between IL-1 signaling and offspring survival, suggest that IL-1A is one of the key ways by which the placenta elicits necessary maternal vascular changes during placentation. The lack of appreciable *IL1R1* in uterine immune cells suggests that fetal-maternal cytokine signaling has become, at least in part, evolutionary decoupled from normal immune function in the uterus. Together, these findings converge on a scenario where placental cytokines have acquired novel developmental functions, and evolved to actively remodel the endometrium to enable continued fetal development.

Our findings suggest that IL-1A helps marginal fetuses survive. Not all fetuses are successful; siblings may vie for capillary access in a manner resembling competitive begging in other species (Parker et al., 2002). Pharmacological dampening of IL-1 signaling showed a significant negative effect on surviving litter size. Fetuses were observed in the process of resorption in both saline and inhibitor-treated pregnancies (**Figure 3d-e**). Indeed, intrauterine resorption is routine in *M. domestica*, and reports tracking litter size over time in *M. domestica* estimate a mean ovulation rate ∼2-3 greater than surviving litter size (Harder et al., 1993). Overproduction and attrition of offspring is common in marsupials, which invest little into offspring during placentation compared to lactation (∼50:1 ratio in *M. domestica*) (Hayssen et al., 1993; Low, 1978; Renfree, 2010, 1983). Abundant production of IL-1 sequestration agents (*IL1R2*) and receptor inhibitors (*IL1RN* - equivalent to anakinra) by maternal cells suggests that the mother actively suppresses IL-1 receptor activation. This could be interpreted as a form of maternal resistance to fetal solicitation: fetuses benefit from signaling that rescues marginal siblings, while the mother curbs excessively large litters (**Figure S3**). Whether this pattern results from sibling competition, parental conflict with (a cooperating cohort of) offspring, or another evolutionary dynamic will require further investigation.

Placental cytokine signaling could conceivably function to induce earlier parturition. Sibling competition has rarely been studied in marsupials *in utero*, but has been proposed during neonatal life. Several marsupials (Virginia opossum and tasmanian devil: Ward, 1998) deliver more offspring than they have nipples, which determines the number they can support and nurse. This creates competition among siblings to reach the nipples first, thought to drive the evolution of traits such as precocious forelimb development (John et al., 2023). IL-1A, IL-6, and IL-17A can induce premature delivery in humans and mice (Gomez-Lopez et al., 2022; Ito et al., 2010; Kaminski et al., 2018; Martínez-García et al., 2011), and on the opposite end, *IL1R2*-activating polymorphisms reduce the risk of preterm birth in humans (Langmia et al., 2016). However, the potential for a lactational bottleneck is weak in *M. domestica*: females have 11-13 nipples (Robinson et al., 1991), a mean litter size at term of around 9 (Harder et al., 1993) (comparable to our colony’s historical modal size), and litter sizes one day before delivery in our study ranged from 7 to 20 with a control group mean of 12. Thus, while effects on parturition timing remain untested, the life history patterns of *M. domestica* suggest that intrauterine brood reduction is more likely the time when competition plays out in this species.

IL-6 inhibition increased fetal biomass by 10% but did not affect survival, implying that placental IL-6 is slightly deleterious to fetal growth. IL-6 is pro-inflammatory via receptor-independent signaling in leukocyte activation, but has anti-inflammatory effects associated with signaling through IL-6R (Scheller et al., 2011), which is widely expressed in opossum uterine cells (**Figure 2c**). In eutherian mammals with epitheliochorial placentas such as pigs, IL-6 is inferred to be anti-inflammatory and aid development of placental villi (Velez et al., 2024), and peripherally, circulating IL-6 drives energy mobilization through lipolysis of fat stores (Kistner et al., 2022). One explanation for our lack of observed growth benefit to placental IL-6 is that it prepares maternal physiology for the metabolic demand of lactation just one day later, with a payoff to neonatal growth realized only after our measurement window. Alternatively, if IL-6 once was a nutrient solicitation signal, it is possible that the mother has evolutionarily modified or dampened the endometrial response to IL-6.

Our results suggest that the classical immunological constraint hypothesis for marsupial gestation is no longer tenable. If inflammation were an alloantigen-triggered or damage-induced form of maternal rejection, we would not expect these signals to originate in fetal tissues, nor to act mainly on non-immune targets. The possibility that this was the ancestral state in early viviparous mammals and secondarily modified has been discussed by Stadtmauer and Wagner (2020). However, a revised constraint hypothesis can be entertained. Many of the cytokines expressed in the syncytium share a common regulatory architecture (Santoso et al., 2020). Unless each one evolved trophoblast expression via independent placenta-specific enhancer elements, they likely cannot vary completely independently, instead evolving as a co-regulated program which as a whole functions positively in placental vascular development. It may not be easy for selection to efficiently prune weakly deleterious signals without affecting other cytokines such as IL-1A which evidently benefit survival. Targeting IL-6 specifically with a drug may over-atomize the phenotype beyond what natural variation could realistically achieve. Identification of the gene-regulatory changes that led to syncytiotrophoblast production of these myriad cytokines in marsupials will be necessary to test this form of immunological constraint.

These are, to us, the most compelling answers to the puzzle of why marsupial fetuses produce signals which seem to limit their own intrauterine growth. Offspring solicitation signals have been extensively modeled in the theoretical literature, frequently under honest signaling models that predict that the only way for solicitation signals to be evolutionarily stable is if they carry associated costs (Grafen, 1990; Kilner and Johnstone, 1997; Mock et al., 2011; Parker et al., 2002). Some aspects of this theory, developed mostly in behavioral contexts, carry over well to marsupial pregnancy whereas others may be misleading. Most importantly, our experiments inhibited receptors rather than ligand production itself. As such, the observed effects of cytokines must have been mediated by the physiological consequences of fetal cytokines on the mother rather than by the metabolic cost of producing signals, which is usually assumed in models of costly begging. Grafen (1990, p. 527) asks that ideas such as “that nestlings beg so noisily because it reduces their growth… not be rejected on the grounds that they are simply absurd”. In the absence of evidence otherwise, we believe it is simply more parsimonious to assume that IL-1’s primary effect is on survival, and that the size differential is driven by more resources becoming available to survivors in reduced litters, while IL-6 is more clearly costly.

The present study has several limitations. First, it was not possible to measure both biomass and gestation length at the same time. *M. domestica* females frequently consume some of their neonates, preventing us from recording both term gestation length and offspring biomass at the same time. Future investigation into effects of placental inflammatory signaling on gestation length is warranted. Furthermore, our study design did not allow us to measure inter-individual variation in biomass or cytokine production among fetuses in the same uterus. If IL-1A promotes redistribution of resources, biomass variance within a litter should be lower in its presence than with inhibition. Evaluation of this variation will be required to rigorously test for sibling competition. Despite these limitations, our experiments allowed a cost of IL-1 and IL-6 signaling, and a positive effect of IL-1 on surviving litter size, to be observed. While our model of uterine capillary development and differential offspring survival remains speculative, it presents testable predictions and a path forward for future investigation into a surprisingly rich and unusual form of fetal-maternal relationship.

Recently, we found that *IL1A* and *IL6* production during the short post-attachment placentation stage of pregnancy is shared between *Monodelphis* and the fat-tailed dunnart *Smithnopsis crassicaudata,* whereas the tammar wallaby *Macropus eugenii* expresses *IL6* but not *IL1A* at appreciable levels (Dudley et al., 2024). The dunnart has an average litter size of 7, similar to *M. domestica*, whereas the wallaby has singleton births (Hayssen et al., 1993). If litter size is regulated by gestational IL-1A in marsupials as our empirical results suggest, this phylogenetic distribution is consistent with a lack of functional utility for fetal IL-1A in a singleton species. Outside of mammals, the viviparous skink *Chalcides chalcides* has a yolk sac placenta analogous to the opossum, a mean litter size of 7.8 (Caputo et al., 2000), and similarly shows expression of IL-1A in its placenta (Paulesu, 1997). Further investigation of the role of placental cytokines in vertebrates with diverse reproductive strategies is warranted.

We can place the opossum’s fetal-maternal signaling into the trajectory of mammalian viviparity more broadly. The mammalian common ancestor’s reproductive mode was likely matrotrophic oviparity, where embryos remain in egg coverings throughout gestation and are nourished by uterine secretions, a state retained in monotremes (Blackburn, 1992). As viviparity evolved, embryos began to hatch internally by secreting proteases that dissolve the proteinaceous shell coat. Continued protease secretion after hatching - conserved across opossums, humans, and rodents (Basanta et al., 2024) - irritates the uterine luminal epithelium (Brosens et al., 2014). Mucosal inflammation leads to angiogenesis and vascular leakage at the site of fetal attachment, as well as neutrophil and monocyte recruitment. Initially, the former was beneficial to fetal development while the latter was detrimental. This provided raw material for natural selection to prune harmful signaling interactions while elaborating useful ones - a process likely central to the evolution of vertebrate placentas (Griffith, 2021; Griffith and Wagner, 2017). Here, we provide evidence that the present evolutionary state in marsupials is not just the result of reductive pruning, nor is it a crude unpruned state as early constraint theories alleged (Cox, 1977; Lillegraven, 1975), but also the product of lineage-specific innovation. Evidence for pruning include maternal expression of IL-1 antagonists IL1RN and IL1R2, and the fact that uterine leukocytes largely lack IL1R1 expression, preventing positive feedback amplifying inflammatory signaling. However, angiogenic fetal signals such as IL-1A and a truncated isoform of VEGFA appear to have acquired developmental functions beyond just byproducts of an immune reaction, and it is possibly through these effects that IL-1A exerts a positive effect on fetal survival. We propose that fetal-maternal communication in the opossum is evolutionarily derived from mucosal inflammation, and has undergone modification both to enhance fetal survival and to protect the mother from over-investment in a single litter.

### Clinical Implications

Meloxicam is regularly administered as a veterinary analgesic to gray short-tailed opossums at low doses of 0.2 mg/kg (Kennedy, 2023). Our failure to demonstrate an effect of meloxicam on local PGE_2_ levels 6 hours post-injection (**Figure S1**) suggests that it is rapidly metabolized. The metabolic rate of meloxicam has been shown to be elevated in several marsupials (brushtail possums, koalas, and ringtail possums) compared to eutherian species like rats and dogs by an order of magnitude, with a half-life on the order of hours (Kimble et al., 2014). The efficacy of meloxicam in marsupials may therefore be overestimated and treatment regimes may be in need of reconsideration. More encouragingly, our observed biomass effect of tocilizumab suggests that it is functionally active in *M. domestica*, unlike the mouse (Lokau et al., 2020). Despite their phylogenetic distance, the opossum exhibits greater IL6RA sequence conservation with the human than does the mouse, including at the 6 amino acid positions known to bind tocilizumab (Lokau et al., 2020): 4 of these residues are identical, while the other 2 are substituted with amino acids with similar charge properties (R→Q and E→K) (**Figure S4**). The opossum may therefore be a more suitable model organism for tocilizumab than murine counterparts.

## Materials and Methods

All experiments and animal care were conducted following ethical protocols approved by the Yale University Institutional Animal Care and Use Committee (nos. 15-11313 and 20-11313).

### Animal Husbandry

*M. domestica* were raised in a breeding colony at Yale University according to established technical protocols (Keyte and Smith, 2008). Male and female animals were housed separately after 3 months of age, at which point female individuals were introduced to the male room for sexual preconditioning. Breeding was attempted after 6 months of age. Non-cycling female opossums were introduced to the male room for 1 day, and subsequently swapped into the used cage of a prospective male partner for 5 days. Afterwards, both individuals were placed into a breeding cage and video recorded to assess the time of copulation. If multiple copulations were observed, the first was used to calibrate 0 dpc.

### Injection Trials

Female opossums were mated as outlined in our husbandry protocol. On day 11.5 of pregnancy, females were assigned to treatment groups of anti-inflammatory (meloxicam, anakinra, or tocilizumab) or a vehicular control of saline. Meloxicam and anakinra were administered at a dose of 1 mg per animal (10 mg/kg for an average 100g female). Tocilizumab was administered at a dose of 3 mg/animal (30 mg/kg) based on the allometric scaling approach of Nair and Jacob (2016) by which this dosage was determined as the equivalent to a 4 mg/kg human dose.

Unless otherwise noted, injections were given on day 11.5, 12.5, and 13.5, and animals were necropsied 6 hours following the last injection (at day 13.75). The ovarian-facing top-third of each uterus was flash-frozen in liquid nitrogen and stored at −80°C for small molecule extraction, and fetuses were collected into pre-dessicated glass vials divided by uterus of origin (two from each animal) for dry weight. Fetuses were desiccated in a 60°C oven for at least 72 hours before weighing.

### Osmotic Pump Implantation

Female opossums were again mated as outlined in our husbandry protocol. On day 12.75 of pregnancy, females were placed in one of two conditions, meloxicam treatment or a vehicular control of saline. Meloxicam and vehicular control were administered via an Alzet osmopump 2001D according to the manufacturer’s recommendations. Pumps were loaded with 200uL of veterinary grade meloxicam (Boehringer Ingelheim Metacam, 5 mg/mL injectable product), equaling a total of 1 mg to match the syringe injections. Given the properties of the pump, meloxicam was released at a constant rate of ∼ 8 μL per hour over a 24-hour period.

Pumps were implanted subcutaneously on the dorsal surface posterior to the scapulae. Animals were anesthetized using controlled isoflurane administration and held in sternal recumbency, and the implantation site was shaved with clippers and washed with povidone-iodine. Pumps were inserted via a mid-scapular incision and the wound closed with wound clips. At 13.75 days from copulation, 24 hours following osmopump implantation, opossums were necropsied and tissue and fetuses were collected as above.

### Quantification of Prostaglandin E_2_

Flash-frozen uterine samples were sent to Cayman Contract Services (Ann Arbor, MI) for commercial quantification of prostaglandin E_2_ using enzyme-linked immunosorbent assay (ELISA) (Cayman 514010 kit; monoclonal antibody #414013). Protein was quantified by bicinchoninic acid assay (BCA; Cayman 701780 kit) and used to normalize PGE_2_ concentrations.

### Statistical Analysis

Opossums have two fully separated uteri, although litters in both are almost always fertilized, and delivered, simultaneously. Given this peculiarity, per-uterus measurements were first obtained and then aggregated to yield per-animal metrics (sum for litter sizes, mean for biomass per fetus across both uteri).

To evaluate the effect of treatment on biomass per fetus and litter size while accounting for the non-independence of measurements from two uteri from the same individual animal, we employed a mixed linear model using the mixedlm() function in the statsmodels python package (v0.14.2) to take advantage of the unique nested structure of our measurements. The model was specified using the formulae “BiomassPerFetus ∼ C(Treatment)” and “Litter Size ∼ C(Treatment)”, where Treatment is a categorical fixed effect. A random intercept was included for each animal. Parameters were estimated using restricted maximum likelihood (REML), the default estimation method for the .fit() procedure in statsmodels, and the same function was used to calculate Wald test *p*-values.

In addition to the mixed model, treatment effects on biomass per fetus and litter size aggregated between both uteri in each animal were examined using two-tailed Welch’s t-tests via the ttest_ind() function in the scipy python package. Both statistical results are reported in order to ensure robustness of findings across approaches. Where appropriate, effect size of treatment was quantified by calculating variance-standardized mean differences (Cohen’s *d*).

As anakinra treatment showed significant effects on both biomass and surviving litter size, analysis of covariance was used to assess whether the treatment effect on these two variables was statistically independent. All per-uterus measurements of fetal biomass and surviving litter size from saline-treated and anakinra-treated individuals where both measurements were available were used for this analysis. The model formula (“Biomass ∼ Litter Size * Treatment”) treated total fetal biomass as the dependent variable, surviving litter size as a covariate, treatment group as a categorical predictor, and contained an interaction term. The model was fit using ordinary least squares via the ols() function and statistical significance was assessed using Type II sums of squares using the anova_lm() function, both in the statsmodels package as above.

### Laser Microdissection and Micro-Bulk RNA Sequencing

Placental-only transcriptomes were captured from laser microdissected samples of fetal tissues directly apposing the uterine lumen in cryosections of 13.5 dpc pregnant uteri. Uteri to be used for this procedure were dissected into phosphate-buffered saline and subsequently immersed Tissue-Tek OCT compound (Sakura Finetek, 4583) in a block mold and flash frozen by exposure to a bath of isopentane surrounded by dry ice, and stored at −80°C. 14-μm sections were prepared from the resulting blocks on a cryomicrotome (Microm HM 500 OM) and placed onto polyethylene naphthalate membrane slides (Leica Microsystems, 11505158) which had been sterilized by ultraviolet irradiation in a biosafety cabinet (Baker SterilGARD, SG-404) for 30 minutes. Sections were processed on a Leica LMD7000 apparatus and excised tissue dropped directly into lysis buffer for processing using a Qiagen RNeasy Micro Kit (74004). RNA libraries were prepared by the Yale Center for Genomic Analysis using a NEBNext Low Input Library Prep Kit (E6420) and sequenced using an Illumina NovaSeq apparatus.

### Transcriptomic Analysis

Single-cell transcriptomes from the fetal-maternal interface of untreated pregnant *M. domestica* at 13.5 dpc and non-pregnant animals were obtained from recently published studies (Basanta et al., 2024; Stadtmauer et al., 2025) (NCBI Gene Expression Omnibus GSE274701 and GSE292958). Differential expression analysis between vascular endothelial cells in 13.5 dpc and non-pregnant animals was conducted using likelihood ratio testing in MAST v3.20 (Finak et al., 2015). For MAST, pregnancy status and biological replicate were modeled as fixed effects in a generalized linear model (method = “bayesglm”), with the number of genes expressed per cell included as a covariate.

Cell-cell interactions were inferred from single-cell data using simple expression thresholding (chinpy v0.0.57, https://gitlab.com/dnjst/chinpy) complemented by statistical analysis of cell type specificity using the LIANA+ (v1.5.1) (Dimitrov et al., 2023) rank_aggregate() function. A custom ligand-receptor ground truth database was adapted from CellPhoneDB v5.0.0 (Troulé et al., 2025) to only include inflammatory cytokines with 1:1 orthologs between human and opossum (archived at https://gitlab.com/dnjst/antiphlogistic_opossum). Only interactions passing expression thresholds 40 counts per million and 15% of cells in the sending and receiving cell populations were retained.

12.5 dpc placental-only transcriptomes (Wang et al., 2014) were retrieved from the NCBI Gene Expression Omnibus (GSE45211) and transcriptomes from whole uteri across the reproductive (Griffith et al., 2017) (NCBI PRJNA393948) and estrus (Griffith et al., 2019) (NCBI PRJNA543903) cycles, as well as opossum endometrial stromal fibroblasts in various culture conditions (Erkenbrack et al., 2018) (NCBI GSE109309), were used for marker gene plotting. Raw reads were aligned to the *M. domestica* genome (Ensembl version 104) and quantified using kallisto v0.45.0 (Bray et al., 2016). Statistical testing for differential expression in bulk data was conducted using the DESeq2 method as implemented in pyDESeq2 v0.5.0 (Muzellec et al., 2023).

## Acknowledgments

Keely-Nicole Wharton and Janelle Avelino oversaw surgical procedures. Rachel Bobo assisted in testing methods for PGE_2_ extraction. Thanks to Mihaela Pavliĉev for the suggestion to test for geometric effects and for support during the writing process. Liam Taylor and Arun Chavan provided insightful comments on the manuscript.

## Competing Interests

Swedish Orphan Biovitrum provided anakinra (Kineret®) and Genentech provided tocilizumab (Actemra®) from programs offering cost-free reagents for basic research. Both companies were provided with experimental design synopses, but neither had any role in data collection, analysis, interpretation, or decision to publish. None of the authors have employment or significant financial interests in either company.

## Author Contributions

G.P.W. and O.W.G. acquired funding and designed pilot experiments. G.P.W., and D.J.S. designed subsequent experiments. G.P.W., J.D.M., O.W.G. and D.J.S. conducted experiments. J.D.M. and D.J.S. performed animal husbandry.

## Funding

This work was supported by the Australian Research Council (#DP200100344 (O.W.G. & G.P.W.), the John Templeton Foundation (#61329) (G.P.W.), the Donnelly Postdoctoral Fellowship of the Yale Institute for Biospheric Studies (O.W.G.), and a grant from the Yale Institute for Biospheric Studies (D.J.S.). D.J.S. was supported by a PhD training fellowship from the National Institutes of Health (T32 GM 007499) (D.J.S.) and postdoctoral appointment at the University of Vienna (D.J.S.). The views expressed here are those of the authors and do not necessarily reflect the views of the funders.

## Data Availability

Measurements are available as a Supplementary Table. Micro-bulk transcriptomes of 13.5 dpc laser microdissected placentae are deposited in the NCBI gene expression omnibus (GEO) at accession GSE302731. Single-cell transcriptomic profiles of the 13.5 dpc fetal-maternal interface were previously published and are available at GSE274701.

---

**Figure S1.**
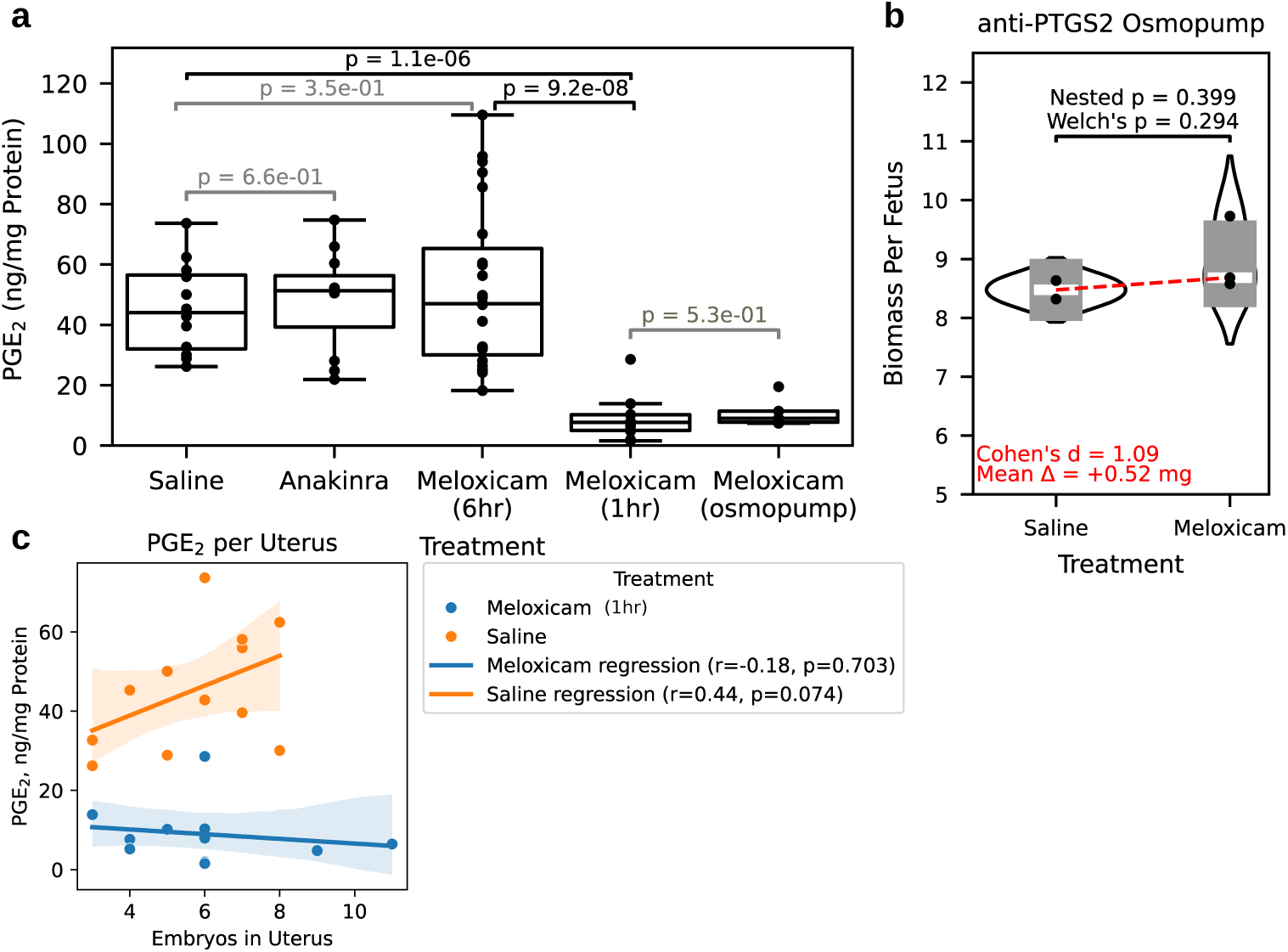
Prostaglandin production under anti-in}flammatory treatment. **a.** Concentrations of PGE_2_ after treatment with meloxicam. **b.** Biomass per fetus remained unchanged after 24 hours of constant infusion via implanted osmopump during days 12.75-13.75 of gestation. **c.** Regression plot of PGE_2_ concentrations versus embryos in uterus shows a positive association in the absence of inhibition.

**Figure S2.**
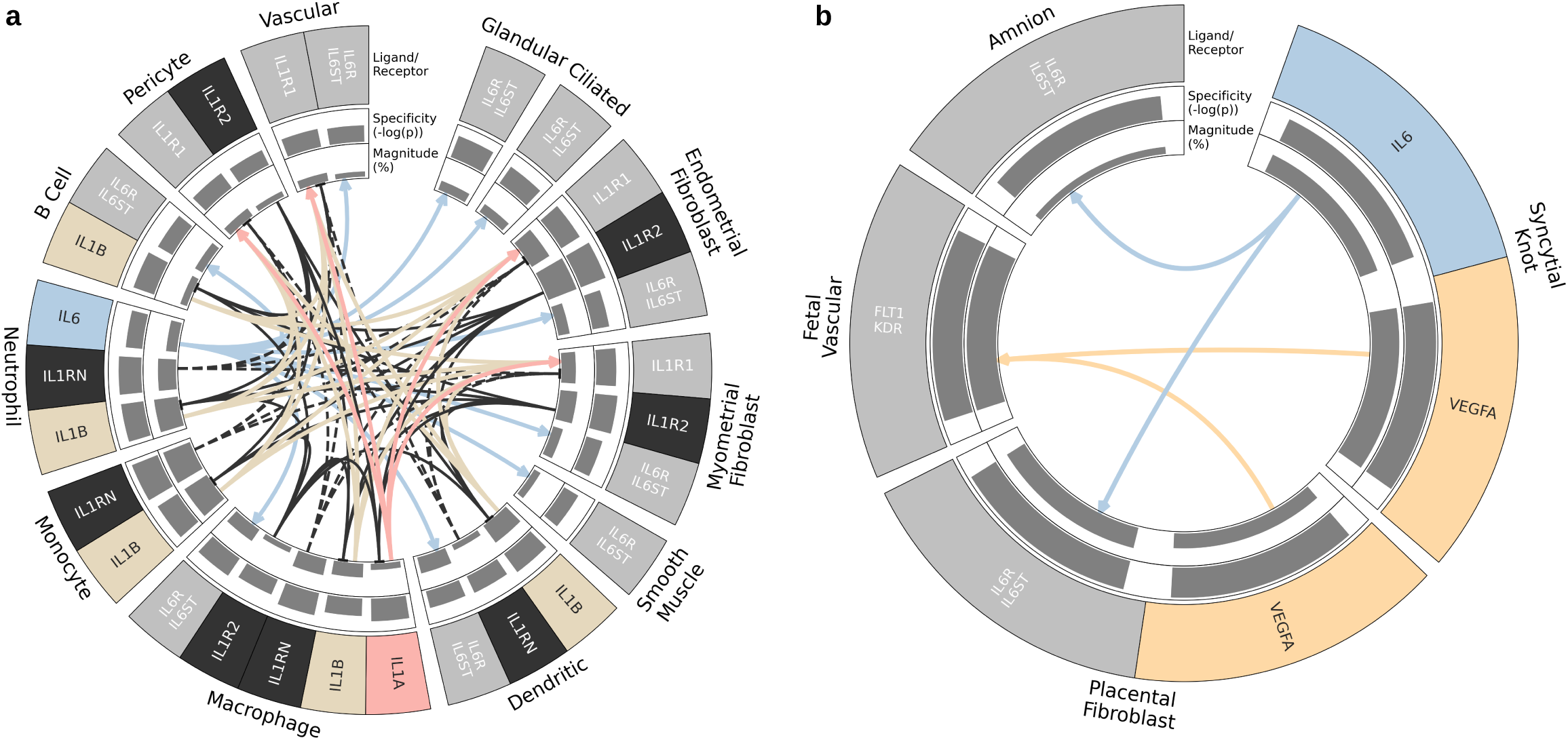
Maternal-maternal (a) and fetal-fetal (b) inflammatory signaling in day 13.5 untreated opossums. Magnitude barplots reflect the percent of cells in the cell type cluster with nonzero expression of the ligand or receptor gene (or least expressed of the two in the case of pairs like PTGS2+PTGES). Specificity barplots reflect the LIANA+ robust rank aggregate p-value scores. Only interactions passing low-expression thresholds for ligand and receptor (15% expression, 50 TPM) are shown.

**Figure S3.**
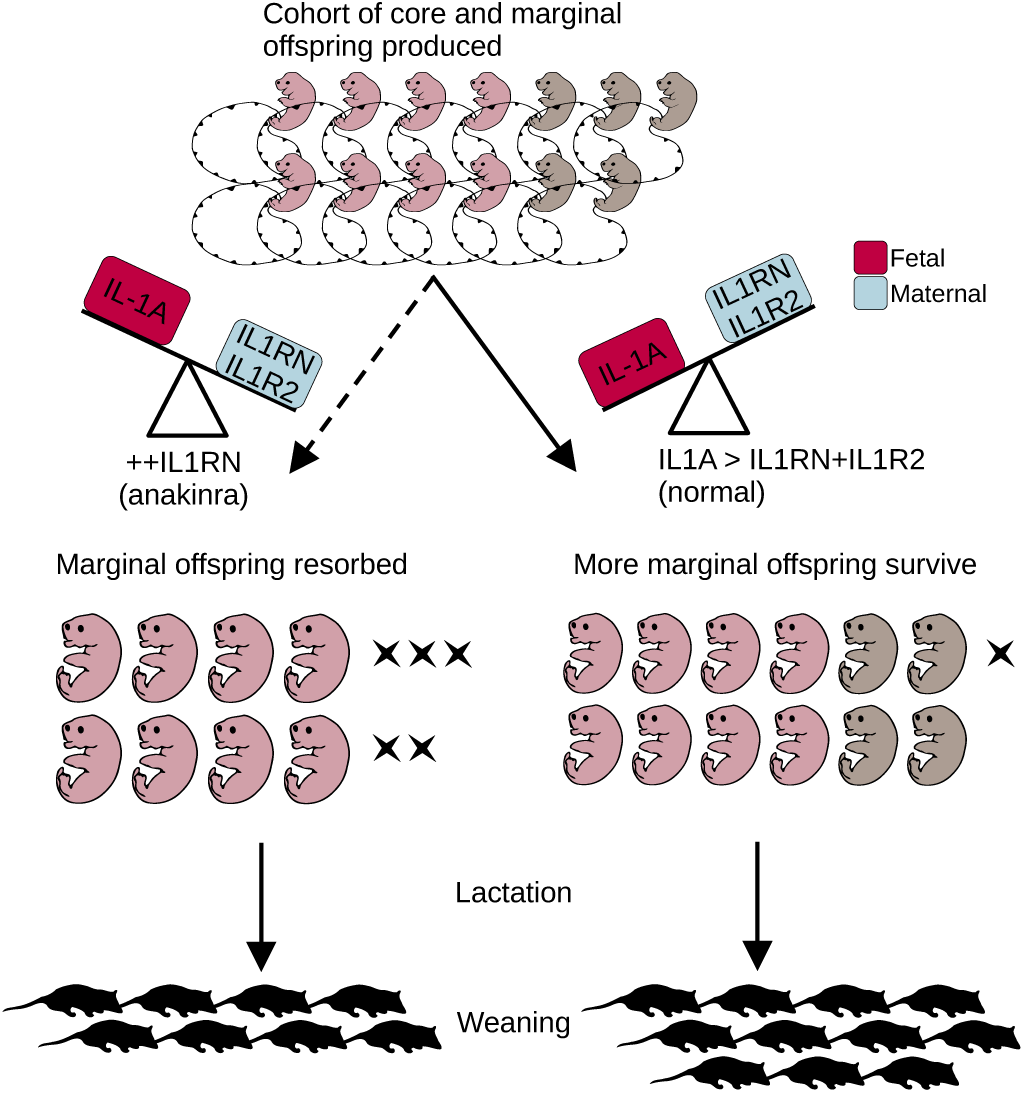
Conceptual model of IL1A’s opposing effects on survival and growth.

**Figure S4.**
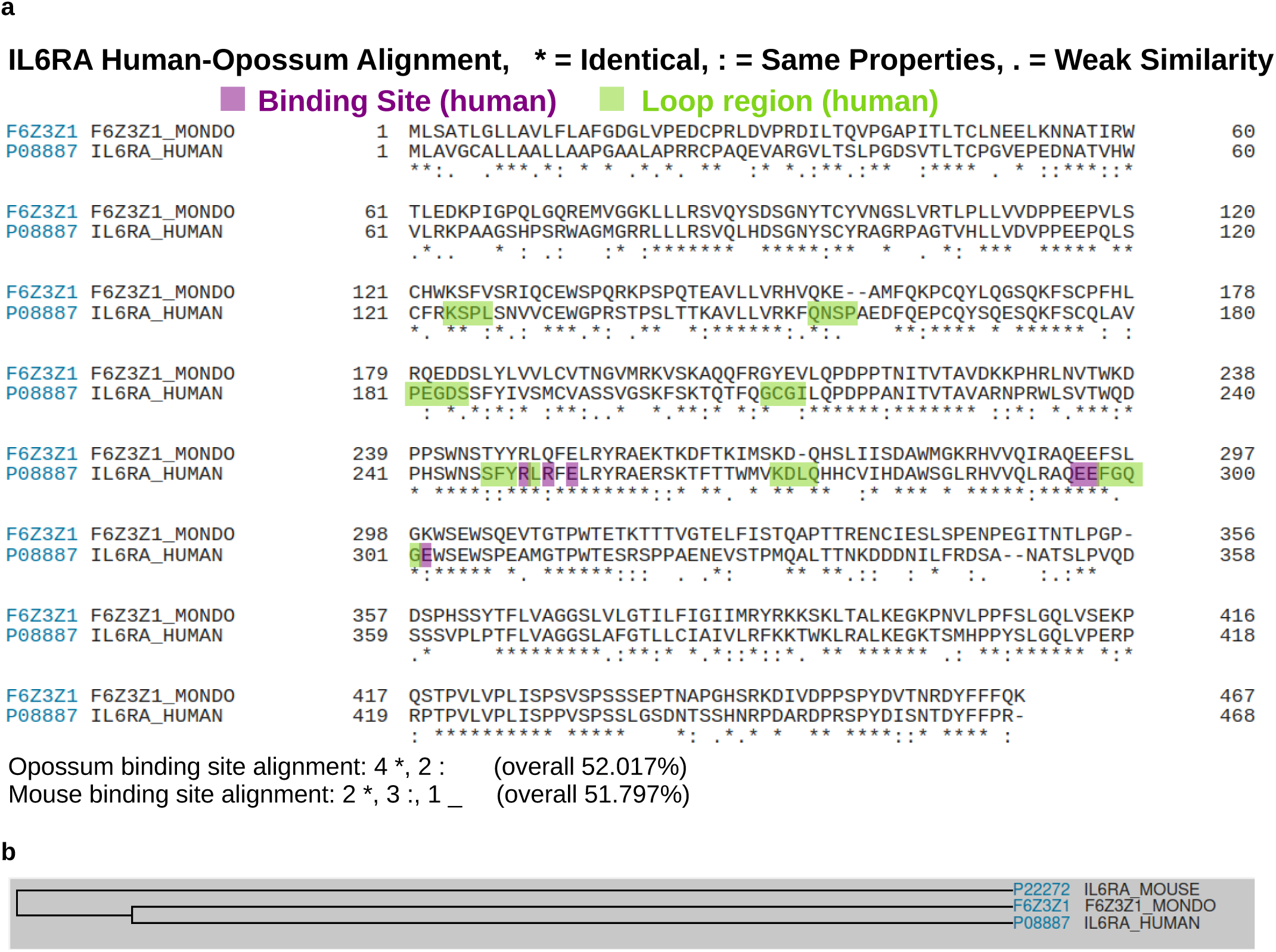
Amino acid alignment between human and opossum IL-6 receptors shows greater similarity to each other than either does with mouse. **a.** Human-opossum alignment of the translated IL6RA peptide. Amino acids constituting the binding site in the human peptide are marked in purple. Substitution codes: * = identical, : = same chemical properties, . = weak similarity, - = insertion/deletion. **b.** NCBI BLAST dendrogram showing greater similarity of opossum and human peptides to each other than to mouse Il6ra, despite the shorter phylogenetic distance between human and mouse.

